# Automatic building of protein atomic models from cryo-EM density maps using residue co-evolution

**DOI:** 10.1101/2020.01.03.893669

**Authors:** Guillaume Bouvier, Benjamin Bardiaux, Riccardo Pellarin, Chiara Rapisarda, Michael Nilges

## Abstract

Electron cryo-microscopy (cryo-EM) has emerged as a powerful method to obtain three-dimensional (3D) structures of macromolecular complexes at atomic or near-atomic resolution. However, *de novo* building of atomic models from near-atomic resolution (3-5 Å) cryo-EM density maps is a challenging task, in particular since poorly resolved side-chain densities hamper sequence assignment by automatic procedures at a lower resolution. Furthermore, segmentation of EM density maps into individual subunits remains a difficult problem when no three-dimensional structures of these subunits exist, or when significant conformational changes occur between the isolated and complexed form of the subunits. To tackle these issues, we have developed a graph-based method to thread most of the C-*α* trace of the protein backbone into the EM density map. The EM density is described as a weighted graph such that the resulting minimum spanning tree encompasses the high-density regions of the map. A pruning algorithm cleans the tree and finds the most probable positions of the C-*α* atoms, using side-chain density when available, as a collection of C-*α* trace fragments. By complementing experimental EM maps with contact predictions from sequence co-evolutionary information, we demonstrate that our approach can correctly segment EM maps into individual subunits and assign amino acids sequence to backbone traces to generate full-atom models.

## Introduction

Getting detailed structural information from cryo-electron microscopy (cryo-EM) is a time-consuming process, which involves sample preparation, data collection, particle picking, and image classification, followed by the reconstruction of the three-dimensional (3D) density map. In the early days of single-particle analysis (SPA) EM, this map used to be called ‘3D structure’. ^1^ However, the key outcome of a cryo-EM study is an atomic model of a macro-molecule or a complex.^2^ In the current paper, we will focus on a method to automatically interpret the final cryo-EM map and generate *de novo*, an atomic model of a protein complex. Therefore, we will ignore the upstream process of obtaining the EM density reconstruction, which will be the input information for the method.

Recent advances in direct electron detectors have led to the structure determination of macromolecular complexes with resolutions as low as 1.6 Å.^3^ The structural features associated with different resolution ranges are the following:

- Below 3.3 Å resolution, the amino acid residue side-chains are visible. This information can be used to unambiguously identify the amino acids and, therefore, to trace the protein sequence into the EM density.
- From 3.3 to 4.5 Å resolution, side-chains are only partially visible. For this ‘shadow’ range, topological reconstruction of the protein is often tedious unless complementary information is available.
- From 4.5 to more than 5 Å resolution, only some secondary structures – mainly helices – are visible. The *β*-strands are no longer separated, rendering the topological reconstruction somewhat impossible.

Since the global resolution of an EM density map is an overall estimation of its information content, the local resolution can significantly vary. ^4^ For example, negatively charged residues that are sensitive to radiation damage or flexible regions have a local resolution that is sensibly lower than the global one. As a consequence, parts of the map may be difficult to interpret, even if the global resolution is sufficient.

Several tools have been proposed to reconstruct protein models from EM density maps. The most common approach is to use structures of the individual components of a complex obtained in isolation by NMR, X-ray crystallography, or comparative modeling. The structures are docked as rigid bodies into the EM map.^5^ These structures can then be further refined, allowing for local flexibility to maximize the correlation with the EM map. This method breaks down when no 3D structures are known. For the highest resolution cases, tools developed for X-ray crystallography are usually applied to cryo-EM, ^6,7^ such as Coot.^8^ Even though automation tools are provided within Coot, the process is largely manual, time-consuming and prone to subjectivity. Furthermore, the sequence assignment is mainly based on side-chain densities to identify the amino acids, which can be impossible in certain parts of the protein.

Other tools have also been developed specifically for EM data. ^9,10^ One of these methods is based on Rosetta^11^ and makes use of the predicted backbone conformation to assign the protein sequence in maps where side-chain density is ambiguous.

One of the first challenges of interpreting *de novo* the EM map of a multi-component complex is the map segmentation and protein identification. Map segmentation is a method that divides the map into sub-maps, each of which corresponding to one unique protein chain of the complex. Other tools exist to find densities corresponding to the asymmetric unit, ^12^ but if the asymmetric unit is composed of distinct protein chains, to further accurately segment remains challenging. To our knowledge, all existing automatic or semi-automatic methods require a previously segmented map.

The goal of our method is to simultaneously segment the EM density map and build an atomic model of all individual protein chains. To do so, we use contacts between amino acids predicted from both evolutionary couplings (EC) and sequence conservation.

## Results

The problem of tracing a chain in the EM density map of a multi-component complex can be divided into several tasks: i) the map needs to be segmented into sub-maps of the individual components; ii) probable locations of amino acid residues need to be identified in the map; iii) these locations need to be assigned to the primary sequences of the components; iv) full atom models of the protein need to be constructed.

The method described in more detail below has four major stages:

1. The cryo-EM map is initially represented as a weighted graph and simplified into a minimum spanning tree (Figure 1(b)) (MST). This tree is pruned until the maximal degree of the graph (i.e., number of edges of the vertex with the greatest number of edges incident to it) is three.
2. The tree is further pruned to remove forks in the tree, and to produce potential fragments of the polypeptide chain. At this stage, the fragments have an arbitrary N-to-C terminal orientation.
3. The fragments are concatenated by making use of evolutionary information in the form of predicted inter-residue contacts. We used the raptorx-ContactMap server^13^ to predict the contacts. The direction of each fragment is determined to maximize the agreement with the predicted contacts. During this stage, the map is segmented automatically, as a consequence of the fragment concatenation procedure. No prior segmentation of the map to individual components is necessary.
4. In the final stage, a full atom model is constructed and refined against the map by standard approaches, Modeller^14^ and the `real space refine` tool^15^ from the Phenix software suite, respectively.

As a test case, we applied our method to the cryo-EM map of the F420-reducing hydrogenase (`EMD-2513`) determined at 3.36 Å resolution.^16^ The map has regions of high and low resolution, correspondingly 3.3 Å and 6.6 Å according to ResMap.^17^ The deposited model (PDB ID `4CI0`), composed of 3 chains of 386, 275 and 281 amino acids respectively, was obtained by a real-space refinement of a former model derived from a previous lower resolution map,^18^ which in turn had been built by the manual fitting of homologous protein structures and subsequent refinement. Regions for which no homolog was available had been manually built using Coot.^8^ In stark contrast, the overall process of our method is automatic and does not require any structural knowledge of a homologous protein. We note that in X-ray crystallography, much higher resolution is required for automated model building. ^19^

**Figure 1:**
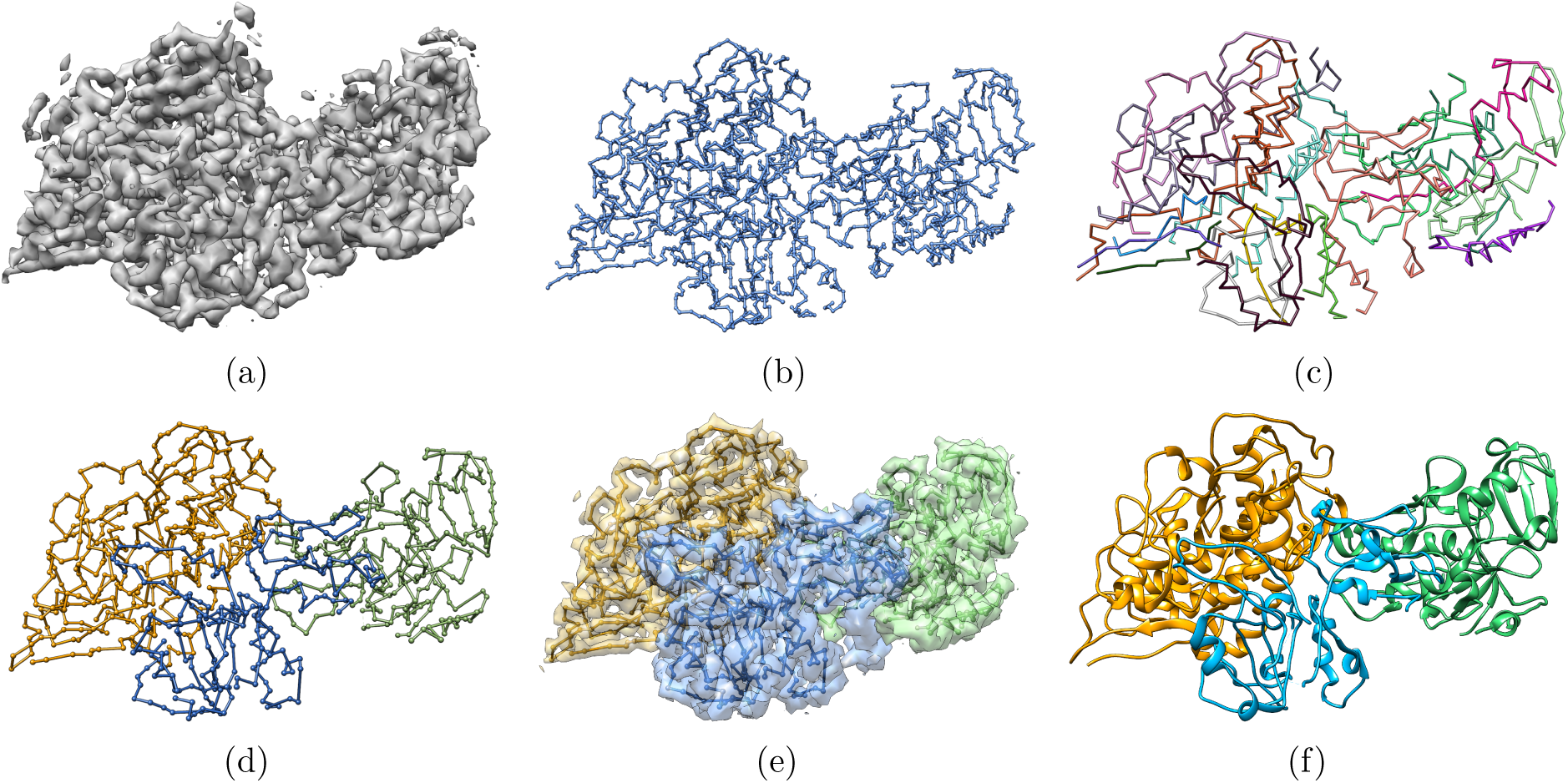
Overview of the automatic reconstruction of an atomic protein structure from an EM density map. (a) Original EM density map used as input of the algorithm. (b) Minimum spanning tree (MSTree) derived from the EM density map. (c) Fragments resulting from the fork pruning procedure of the MSTree. (d) Chains built from the fragments. Fragments were merged in a way to maximize the overlap with predicted contacts. Chains A, B and C were detected automatically without any prior segmentation of the EM density map. (e) Segmented map built from the threading of the 3 chains depicted in d. (f) Full atom model derived from the C*α* tracing in d) with Modeller and the `real space refine` tool from the Phenix software suite.

The inputs of the algorithm are the map of the asymmetric unit consisting of a heterotrimeric protein complex (Figure 1(a)), the primary sequences of the three protein subunits, and the predicted contacts for each subunit. RaptorX provides a score, given as a probability, for each predicted contact. We used a cutoff of 0.5 on the prediction score to discard irrelevant contacts, ending up with 658, 343, and 563 predicted contacts for chains A, B, and C, respectively. The average number of contacts per residue is consequently 1.7, 1.2, and 2.0 for the 3 chains respectively.

For our test case, the algorithm found 18 fragments (Figure 1(c)). The fragments were placed in the density with an average accuracy of 1.2 Å, computed as C-*α* RMSD to the reference structure, `4CI0`.

The generated fragments fully cover the three chains of the reference structure `4CI0` (Figure 2). In general, the fragments tend to correspond to clusters of secondary structure elements, and the ends of the fragments lie at the edges of secondary structures.

**Figure 2:**
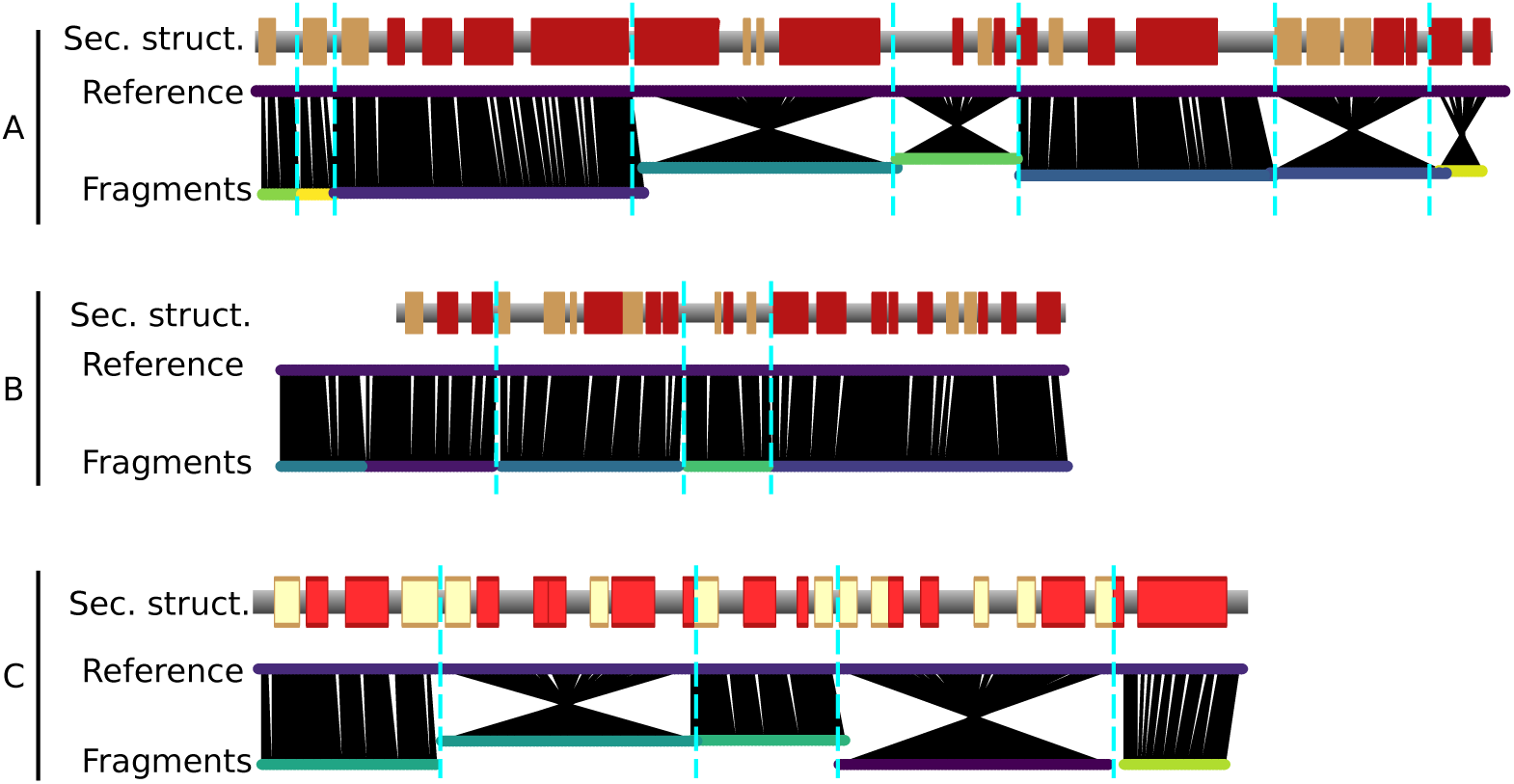
Fragments resulting from the map threading (bottom lines) and aligned on the 3 chains of the reference model `4CI0` (upper lines). The red and yellow boxes indicate *α*-helices and *β*-strands, respectively in the reference model. The black lines indicate the nearest C*α* atom in space from the fragment to the reference model. Crossed line bundles indicate fragments that have been reversed with respect to the originally predicted direction of the graph.

As mentioned, the identified fragments do not contain any information on the direction of the polypeptide chain. Hence, both directions of the fragments have to be tested to assign the protein sequence, thus considerably increasing the combinatorial complexity of the problem, as twice the number of fragments have to be considered. Chain direction inversions are clearly apparent in Figure 2 as ‘crossed’ features.

The resulting identified chains are shown in Figure 1(d). In the example, the proposed segmentation fits extremely well the chains of the reference structure (PDB `4CI0`). In contrast to other tools developed for *de novo* protein structure determination from EM maps, which consider segmented maps for individual chains as a prerequisite, ^2,11^ as mentioned earlier, the current approach performs map segmentation and tracing of the chain simultaneously.

We compare the final backbone contact maps with the predicted contact maps in Figure 3. The overlap score is defined as the fraction of predicted contacts also observed in the model. To get a local overlap, we define a window of 11 consecutive residues and slide it along the sequence. We compute the overlap score of the 11-mer with the target structure for each residue position (Figure 3).

**Figure 3:**
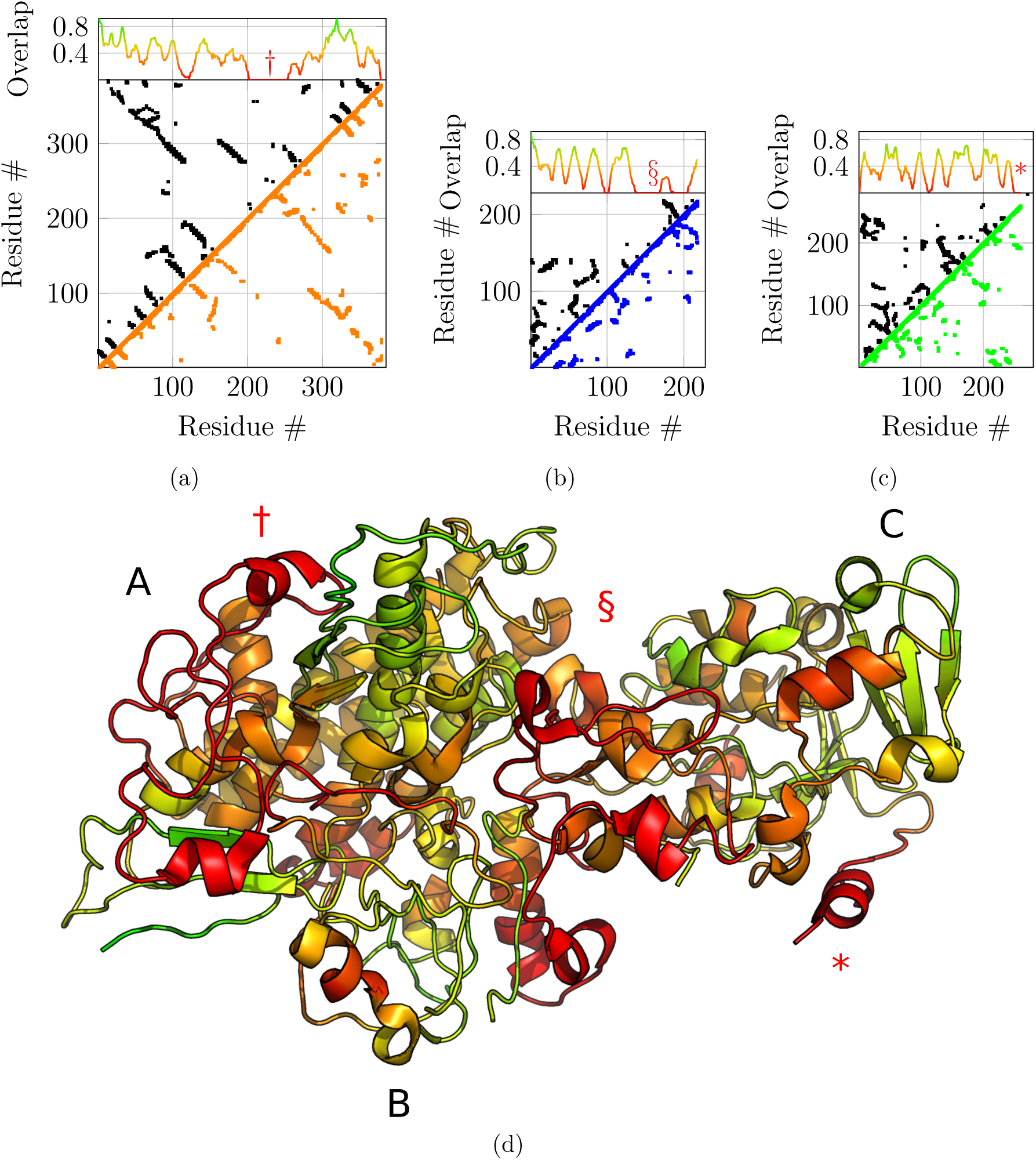
Predicted contact maps – upper diagonals in black – with a probability cutoff of 0.5, compared with the final contact maps of chain A, B and C ((a), (b) and (c) respectively) resulting from fragment merging – lower diagonal in orange, blue and green respectively – with a distance cutoff of 8 Å. The upper projection gives the overlap ratio between the contact map of the model and the predicted contact map. An overlap of 1 means that all predicted contacts are satisfied by the model. The overlap is computed for sliding fragments of 11 residues along the sequence. (d) Projection of the contact map overlap on the generated model, colored from red (overlap 0) to green (overlap 1). Three regions, highlighted by symbols †, § and * for chains A, B and C respectively, are regions with very sparse predicted contacts and therefore very low overlap scores, indicating that the sequence assignment for these regions on the final model could be wrong.

In order to assess the accuracy of the sequence assignment based on the predicted contact map, we color-coded the computed structure according to the overlap score (Figure 3(d)).

Next, we compared the obtained model to the original structure `4CI0` (Figure 4). Deviations of C*α* positions from the reference model are plotted in figure 4(a), 4(b) and 4(c) for chains A, B and C respectively. Chains A and C are accurately placed with an average deviation of 2.75 Å and 2.31 Å, respectively. Globally, regions with higher deviations are contiguously found around residues 250 for chain A – noted with symbol † in Figure 4(a) – and for the C-terminus of chain C only – noted * in Figure 4(c) – and are well delimited in the structure (Figure 4(d)). These regions correspond to the regions where the predicted contacts are sparser than the other parts, highlighted above with symbols † and *. Chain B is less accurately placed with an average deviation of 4.17 Å. The deviation is essentially due to residues 150 to 200 – noted § in Figure 4(b). This deviation is explained by the low overlap score obtained for that region (§ marker in Figure 3(b)) corresponding to the region with very sparse predicted contacts, combined with the local low resolution of the cryo-EM map.

**Figure 4:**
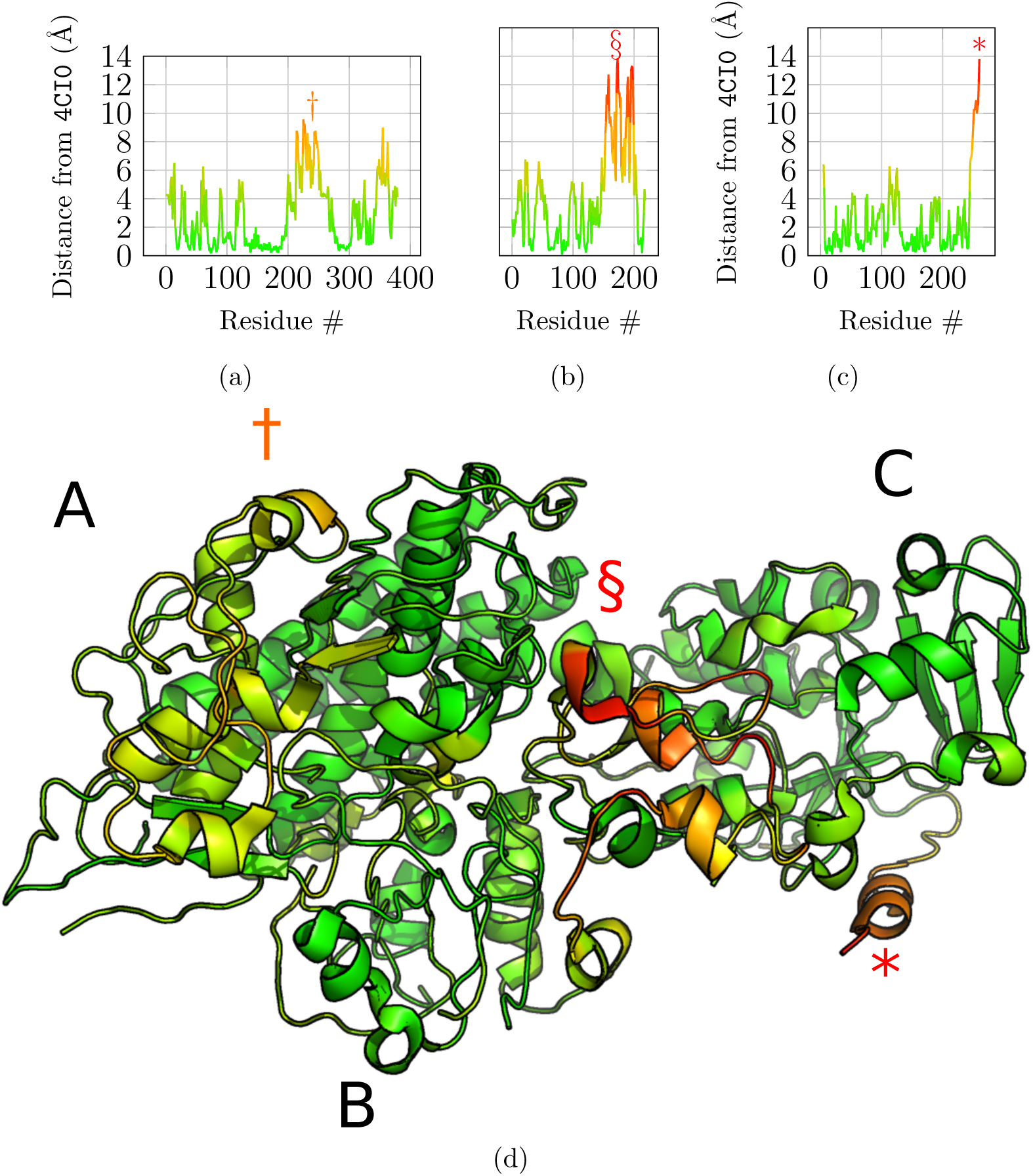
Structural alignment and comparison with the reference model `4CI0`. Graphs a, b and c shows the deviations of the final model for each of the C*α* atoms from `4CI0` aligned to the model, for chains A, B and C respectively. (d) Final model with C*α* deviations projected onto the structure using the same colormap as for graphs a, b and c. Symbols †, § and * highlight regions in the structure with high deviations from the reference structure for chains A, B and C respectively.

We computed the local correlation per amino acid residue to assess the fit to the cryo-EM data of the refined reconstructed model (Figure 5). For this, we used the Segment Manders’ Overlap Coefficient (SMOC) as described in^20^ with a sliding window of 11 residues, similarly to the calculation of the overlap score. Qualitatively, the protein regions that diverged in C*α* positions from the reference structure are also poorly fitted in the density, especially for the region noted § for chain B (Figure 5(b)). Therefore, the local correlation of the obtained model with the EM density map can be used to identify regions to carefully analyze and manually re-model when no structural data is available. An incorrect register of the sequence can explain those results due to poorly predicted contacts for this region or to a poor local resolution of the density map.

**Figure 5:**
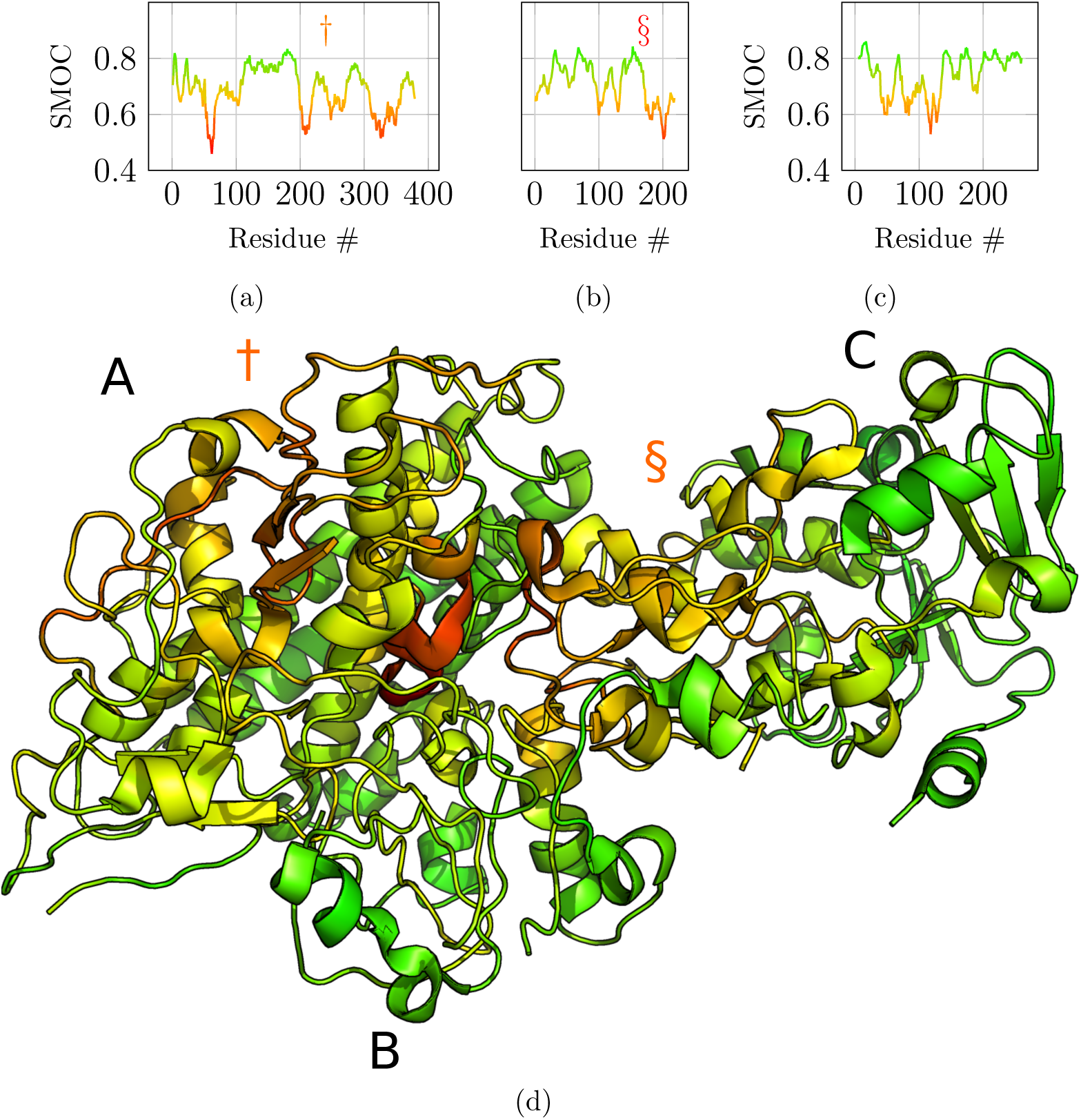
Segment Manders’ Overlap Coefficient (SMOC) of the refined model against the EM density map `EMD-2513` used as input for the atomic scale reconstruction method. The colormap used highlights poorly fitted regions in red. SMOC profiles along the protein sequence are depicted for chain A, B and C in panels a, b and c respectively. (d) The corresponding SMOC values are projected on the obtained full atom model.

The method above was adapted to be used with manually traced backbone. We recently applied this modified methodology to determine the structure of the wedge complex of the bacterial Type 6 Secretion System (T6SS) baseplate.^21^ The wedge complex is mainly composed of three proteins: TssF, TssG and TssK. The cryo-EM density map (EMD-0008) is organized into three lobes, with distinct resolutions. Two of the lobes were occupied by two TssK trimers. In these regions the resolution and structural prior information was enough to build accurate atomistic models. In contrast, the resolution of the lobe corresponding to TssG and TssF was between 4.3 and 8 Å. Since only weak homology information was available for TssG and TssF, the structure of the proteins was determined *de novo* using a semi-automatic iterative pipeline, fully described in. ^21^

In this application, we did not employ the automatic backbone tracing of steps 1. and 2., described above, since the resolution was too low for an automatic tracing procedure. The TssFG map was segmented using Segger, ^22^ which identified densities corresponding to TssG and two TssF subunits arranged in a pseudo-C2 symmetry. The C-*α* backbone structure was traced manually based on the densities of the three segment using Coot. The tentative sequence registering was obtained by aligning the bulky amino acids and using secondary structure prediction. The manually traced backbone was highly inconsistent with the predicted contacts (Fig. 6A and F). To resolve these inconsistencies, we iteratively applied two steps: first, we fragmented the backbone based on the alignment of the contact map of the model on the predicted contact map. The alignment identified gaps with more than four residues, that we used to enforce the fragmentation position. Second, we concatenated the fragments following stage 3 discussed above. For TssF, we needed six iterations of fragmentation and concatenation to obtain a model fulfilling most of the predicted contacts (Fig. 6B and G).

**Figure 6:**
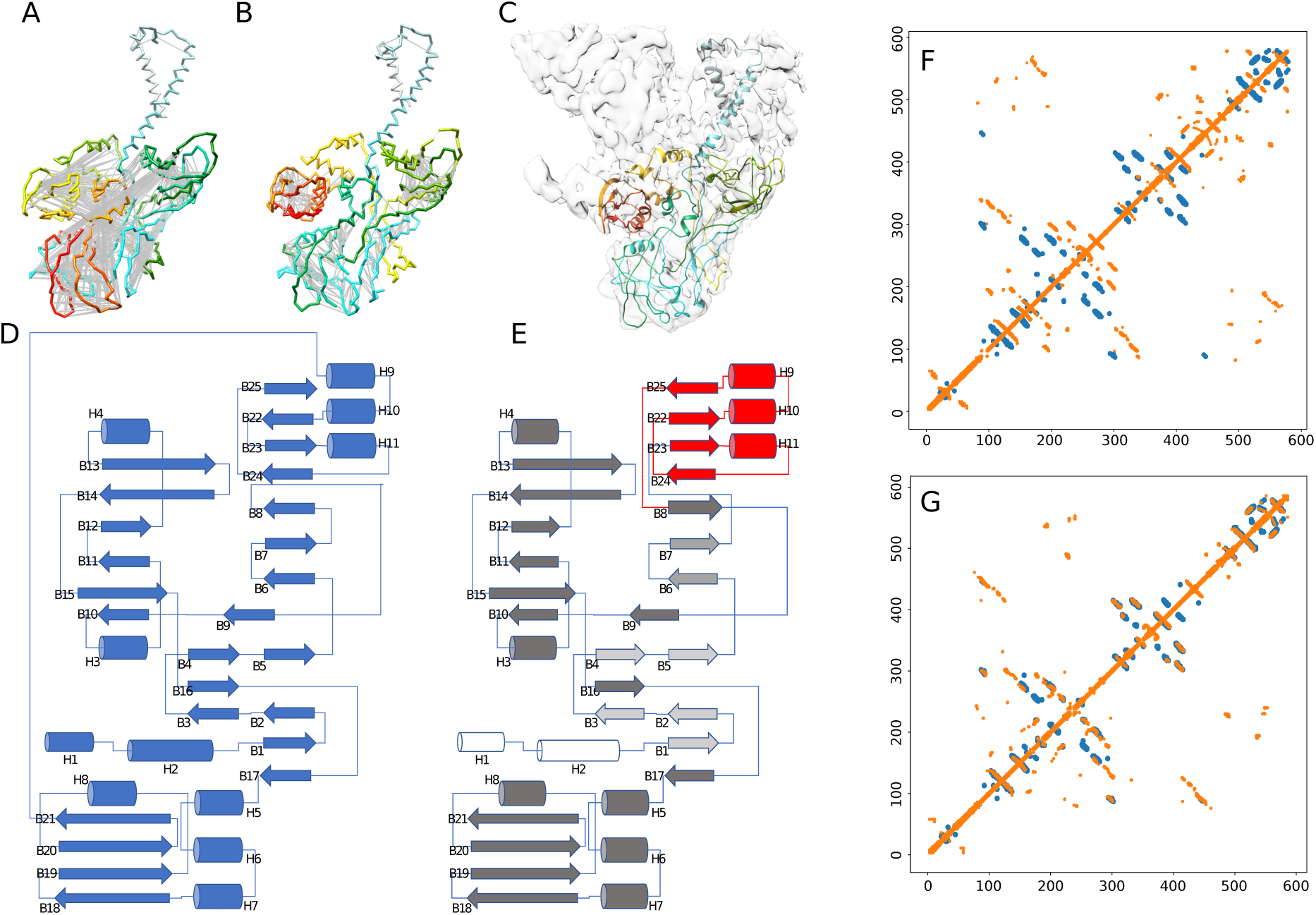
Modified methodology to determine the structure of the wedge complex of the bacterial Type 6 Secretion System (T6SS) baseplate. A) Initial model manually built in the EM density map. B) Fixed model using predicted contacts. Grey lines corresponds to contacts predicted by RaptorX, mostly unsatisfied in the initial model. The rainbow color of the backbone report the residue index, where N-terminal residues are blue, and C-terminal residues are red. C) Final atomic model docked in the EM map to the wedge complex. D) Topology of the initial model depicted in A. E) Topology of the fixed model depicted in B. The red part highlights the major topology changes made by the method (fragmentation/concatenation) The gray scale encodes the sequence offset introduced by the method: white secondary structure elements have no offsets with respect to corresponding secondary structure of the initial topology, light gray elements have an offset of about 10 residues, dark gray have an offset of about 100 residues. F) Contacts of the initial model (orange) aligned on the predicted contacts (blue). G) Contacts of the final model (orange) aligned on the predicted contacts (blue).

With respect to the topology of the manually traced structure (Fig. 6D), the method reoriented three beta-strands, two helices, and changed the connectivity of four pairs of secondary structure elements, localized in the center of the sequence (red secondary structure elements in Fig. 6E). These changes had a huge impact on the topology, considering that two third of the sequence was offset by about 100 residues (dark-grey secondary structure elements in Fig. 6E).

We were able to build an atomistic model of TssF and TssG (not shown) components of the complex, in which most of the sequence of the proteins could be assigned to the cryo-EM density and secondary structure elements could be identified (Fig. 6C). This same structure was later confirmed by other authors using higher resolution cryo-EM maps,^23^ with C*α* RMSD of 3.6Å and 1.8Å for TssF and TssG respectively, confirming the accuracy of the method.

## Discussion

In the first test case, our automatic reconstruction method results in a model with an average accuracy of 2.4 Å computed as the C-*alpha* RMSD to the deposited structure. The map was automatically segmented in the process of sequence assignment. To our knowledge, our method is the only one able to segment the EM density map, in individual protein chains, with only sparse and noisy contacts from residue co-evolution, and, remarkably, without the use of 3D structures of homologous proteins.

A recent strategy, MAINMAST,^9^ makes also use of the MSTree in the early stages of the modeling. However, the tree is refined using a Tabu-search algorithm, and the algorithm is applied to already segmented maps, due to the complexity of the combinatorial problem with larger systems containing multiple subunits. Furthermore, the MAINMAST approach employs a threading approach to register the target sequence on the traced chain by evaluating the fit of the amino acid sequence of the protein to a path in a tree. Our method uses a different approach as the sequence is registered using an alignment procedure between the contact map of the protein model built with the predicted contact map using co-evolution.

Obviously, in some cases, manual intervention will eventually be necessary. The overlap score (Figure 3) helps to identify regions of the protein that might be problematic. The test case illustrates how the overlap function correlates with divergence from the reference structure `4CI0` (Figure 4). Hence, this score provides an objective metric to identify troublesome regions where local modification of the 3D structure might be necessary, or place fragments and assign their sequence by hand to maximize the overlap with the map. Furthermore, when the minimum spanning tree and pruning procedures lead to incorrect topology, the overlap score should pinpoint the misleading regions, thus allowing to fix the error if possible.

The fragment placing and merging procedures, and the sequence assignment are two independent processes in the workflow. Therefore, the method is very flexible as any fragment building method can be coupled with the merging procedure based on the predicted contact maps. In particular, manual tracing of fragments can be easily coupled with our sequence assignment method in cases where the map is of insufficient resolution or quality to place atoms or fragments automatically. In this spirit, we have used the fragment merging and sequence assignment functionalities to solve the unknown structure of the TssF protein, a subunit of the T6SS baseplate, using 5 Å resolution map in combination with manual tracing.^21^

In the current implementation, inter-protein contacts are not yet taken into account but the method can be readily adapted. Alternatively, predicted inter-subunit contacts can still be used with the existing implementation to validate the final model.

The major limitation of the method is at present the collection of reliable contacts form residue co-evolution. Indeed, a sufficient number of homologous sequences is required to predict contacts with an acceptable degree of reliability.^24^ This limitation will likely be of less importance in the future with the availability of contact prediction methods that require fewer sequences. ^25^

## Materials and Methods

An EM density map is stored as a cubic grid, where each grid point encodes the local electron density. The map of the test case is the asymmetric unit (AU) of the EMD-2513. The map of the AU, composed of 3 protein chains, as been obtained using UCSF Chimera’s ‘zone tool’^26^ and results in a grid of 80 × 74 × 76 grid points with a spacing of 1.32 Å.

### Tree representation of an EM map and minimum spanning tree

The first step of our method is to describe the EM map as a weighted graph *G*(*V, E*). The vertices *v* ∈ *V* are the grid points of the EM map, and the edges *e* ∈ *E* are the 26 direct neighbors in the grid. The weight *w*_*i,j*_ of the edge connecting vertices *v*_*i*_ and *v*_*j*_ is defined as the inverse of the product of the electron density of the corresponding grid points noted as *d*_*i*_ and *d*_*j*_, respectively, *w*_*i,j*_ = (*d*_*i*_ × *d*_*j*_)^−1^.

If we assume that the protein backbone is visible in the EM density as a high-density feature, finding the chain trace should correspond to finding the path of minimal weight in such a graph. Therefore, a useful way to simplify the graph is to build the minimum spanning tree (MSTree) of the graph. By definition, the MSTree is a subset of edges of the graph that connects all the vertices without any cycles and with the minimal total weight. Therefore the MSTree should encompass the correct backbone threading in the EM map. We use the Kruskal algorithm^27^ to build this tree from EM density graph *G*.

In a third step, the tree is pruned, by removing portions of the graph until the maximal degree of the graph, i.e. the maximal number of edges per vertex, is three. This simple pruning rule allows us to obtain a potential candidate for the threading of the C*α* trace through the density since C*α* atoms are covalently bonded to three non-hydrogen atoms at most. The resulting tree is depicted in Figure 1(b). We note that the MSTree is similar to the result of the skeletonization procedure often used in *de novo* model building from density maps to guide manual threading of the C*α* trace.^8^

The final goal of the graph algorithm is to obtain the longest fragments describing the best possible partial C*α* threading of the protein backbone. Therefore, as the spacing between the nodes of the graph is still equal to the spacing of the input EM density map (1.32 Å in our case), one must place C*α* atoms as accurately as possible. Starting from remaining vertices of degree three, which are likely C*α* atoms with visible density for the side-chains (or at least C*β* atoms), vertices coordinates are refined to get as many vertices separated by 3.8Å as possible – i.e., the average distance between two consecutive C*α* carbons – (Figure 7(a)).

**Figure 7:**
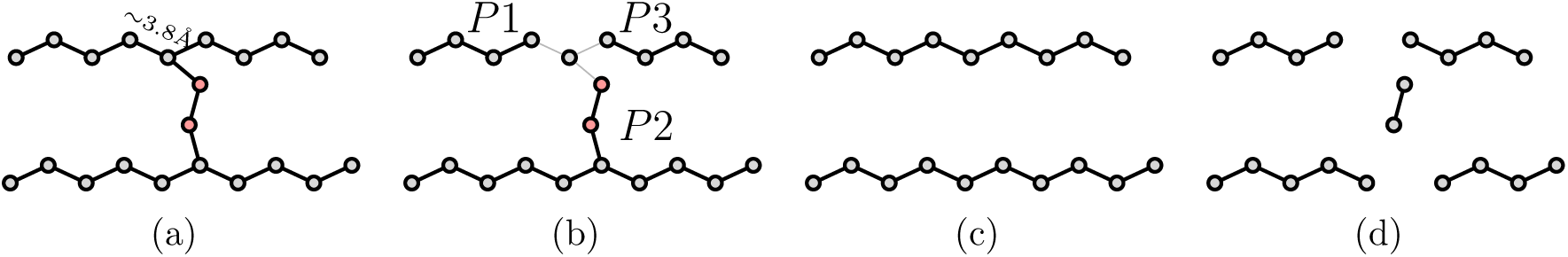
Fork pruning procedure. (a) Regularization of the nodes of the MSTree to be spaced by 3.8Å. The anchoring C*α* defining the reference position to regularize the positions are defined by nodes of degree 3. These nodes are involved in forks (in red) that must be pruned to recover possible backbone threading. (b) Fork pruning procedure: when a fork is detected the edges of the fork are temporarily removed, which creates 3 possible path graphs (*P*1, *P*2 and *P*3). (c) The shortest path graph (*P*2) is removed to solve the fork. However a pruning threshold (*T*_*p*_) is defined to avoid the removal of large fragments. If the length of the path graph is lower than this threshold the path graph is removed and the fork is solved. (d) In the case that the length of the shortest path graph is higher than *T*_*p*_ the fork nodes are simply deleted and more fragments are generated.

The resulting graph can still contain forks. We thus developed an additional pruning procedure to suppress remaining forks and to get unbranched fragments that approximate the backbone threading in the EM density map (Figure 7).

### Fragment assembly and map segmentation

The fragments then need to be assembled to build the protein structure, and the sequence of the chain (or several chains) has to be assigned to the detected C*α* positions. For this purpose, we use the predicted contact map of the chain. This part of the procedure is based on the `map align` algorithm developed by Ovchinnikov et al. ^28^ The `map align` algorithm takes two contact maps and returns an alignment that tends to maximize the number of overlapping contacts while minimizing the number of gaps. Along with the alignment, `map align` gives a score for the quality of the alignment.

However, this algorithm was designed to match predicted contacts with contact patterns of known protein structures and not for *de novo* building of protein chains from fragments. In order to apply the map alignment tool to this purpose, we developed an algorithm that iteratively concatenates the fragments built to maximize the alignment score. For an arbitrarily chosen chain of the complex, the algorithm finds the fragment and its direction that maximizes the `map align` score with the predicted contact map. For this current fragment, the algorithm considers all its neighbors and calculates their tentative `map align` scores. The newly calculated tentative `map align` scores are compared to the current ones, and the new current fragment is chosen as the fragment with the highest score and so on. The algorithm stops after a chosen number of iterations or when all combinations have been tested.

Dynamic programming allows us to find the subset of fragments and their associated directions by maximizing the overlap with the predicted contact map for each chain. Once the first chain has been built, the full set of fragments is reused to get the most probable second chain, and so on for the remaining chains. Then the chains are sorted by alignment scores. The fragments used to build the chain with the highest score are removed from the pool of fragments, and this chain is considered as assigned. Then, the second-highest scoring chain is built in the same way but from the new subset of fragments, preventing overlap of the newly built chain with the previous one, and reducing the combinatorial complexity of the problem simultaneously. This is repeated for each chain until the last chain is built. In this way, the algorithm proposes a segmentation of the EM map based on the predicted contacts.

To compute the contact maps derived from the fragments, we use a distance cutoff of 8 Å between C*α* atoms. Furthermore, the RaptorX contact prediction server provides a score, given as a probability, along with each contact identified. For the current test case, we used a probability threshold of 0.5 to derive the predicted contact maps.

Lastly, the final contact map alignments allow registering the sequence of the subunits onto the C*α* traces as it provides the correspondence between a bead position and the corresponding amino acid position in the sequence used to predict the contacts.

### Building the full atomic model

In the final step, we use Modeller^14^ to obtain a full-atom model of the structure. The obtained model is further refined in the EM map (Figure 1(f)) using the `real space refine` tool^15^ from the Phenix suite. During the refinement, secondary structures are optimized using an optional feature in Phenix that annotates secondary structure based on C*α* positions.

## Acknowledgement

We would like to thank Rémi Fronzes (IECB, Bordeaux) for his contribution to the segmentation of the T6SS baseplate cryo-EM density map.

